# Comparison of Two Budding Yeast Disome Model Systems: Similarities, Difference, and Conflict

**DOI:** 10.1101/2022.12.11.519961

**Authors:** Peter J. Vinton, Rachel E. Langston

**Author notes:** Corresponding Author: Peter J. Vinton.

## Abstract

Here we define chromosome instability as the propensity of error-prone DNA repair and maintenance to generate chromosomal alterations known as gross chromosomal rearrangements (GCR), which can be found in a variety of forms in a variety of diseased cells. Insights and study of GCRs and chromosome instability can be gained through experimentation using a disome system (a haploid strain with an extra copy of one chromosome). Chromosome instability has previously been investigated and identified in a budding yeast ChrVII disome model. Here we extend and compare the study of chromosome instability using a similar ChrV disome system. As with the ChrVII disome system, cells containing unstable chromosomes form a distinctive “sectored” colony phenotype and through the use of genetic markers, we also find evidence of allelic recombination and chromosome loss. We also found the same DNA integrity pathways suppress chromosome instability and that unstable chromosomes are not generated through homologous recombination (HR) or non-homologous end-joining (NHEJ), similar to the ChrVII system. But in contrast and more interestingly, we did not detect any altered ChrV sizes, which conflicts with a previous ChrVII disome study where it was thought that unstable chromosomes often resulted in altered sizes. We also discovered a distinct increase in frequency of instability in the ChrV system compared to the ChrVII system.

## INTRODUCTION

There is a strong connection between chromosome instability, GCRs, and various cancers, which can clearly be seen in the spectral karyotyping of chromosomes from chronic myeloid leukemia, bone cancer, and prostate cancer cells in which regions of different chromosomes have fused together and/or where regions of chromosomes have been lost (Calabrese et al., 2000; Pan et al., 1999; Rogatto et al., 1999). One of the initial steps in generating an unstable chromosome is the occurrence of DNA errors (e.g. single and double strand DNA breaks, base mismatches, pyrimidine dimers, etc.) which often arise stochastically during DNA replication and repair (Mills et al., 2003). DNA errors are usually faithfully repaired, but at times when repair is faulty, GCRs can occur (Chen & Kolodner, 1999). GCRs assume a variety of forms including deletions, fusions, translocations, allelic recombinants and chromosome loss as well as unstable chromosomes with often unknown structures. Unstable chromosomes are particularly intriguing as they may cause cycles of instability and have been identified as a common intermediate in studies of both *Saccharomyces cerevisiae* (Admire et al., 2006; Cobb et al., 2005; Myung & Kolodner, 2000; Vasan et al., 2014) and mammalian cells (Breger et al., 2005; Hill & Golic, 2015; Miller et al., 2015). The transient nature of unstable chromosomes can cause different growth rates in the progeny of the cell which originally contained unstable chromosome(s) due to different levels of DNA repair, types of DNA repair, amount gene loss etc., necessitating the development of systems for quantitative and systematic analysis.

Investigating chromosome instability and GCRs in multiploid cellular models is difficult due to their inherent complexities and redundancies. On the other hand, investigating instability in a simpler haploid system is limited due to the fact that GCRs frequently result in the loss of essential genes, lethal to the cells being studied. To overcome this paradox, a budding yeast ChrVII disome system is an appropriate model (developed by T. Formosa, modified by T. Weinert, no associated publication). This system employs haploid cells with an extra copy of one of the chromosomes. A disome system reduces chromosome complexity yet allows the study of complex chromosomal rearrangements. The ChrVII disome system contains discernable genetic markers in the extra non-essential chromosome that allows detection and characterization of GCRs resulting from chromosome instability (Fig. S1). Here we further study chromosome instability by experimenting with a similarly constructed ChrV disome system (developed by the Alison Adams and Weinert Labs) (Fig. 1). Trends in instability remain similar, such as unstable sectored colony shape, genes affecting instability, and varied genotypes of cells within a sectored colony, but there are also differences between Southern Blot detection of altered chromosome sizes and frequencies of instabilities between the two disome systems.

**Figure 1.**
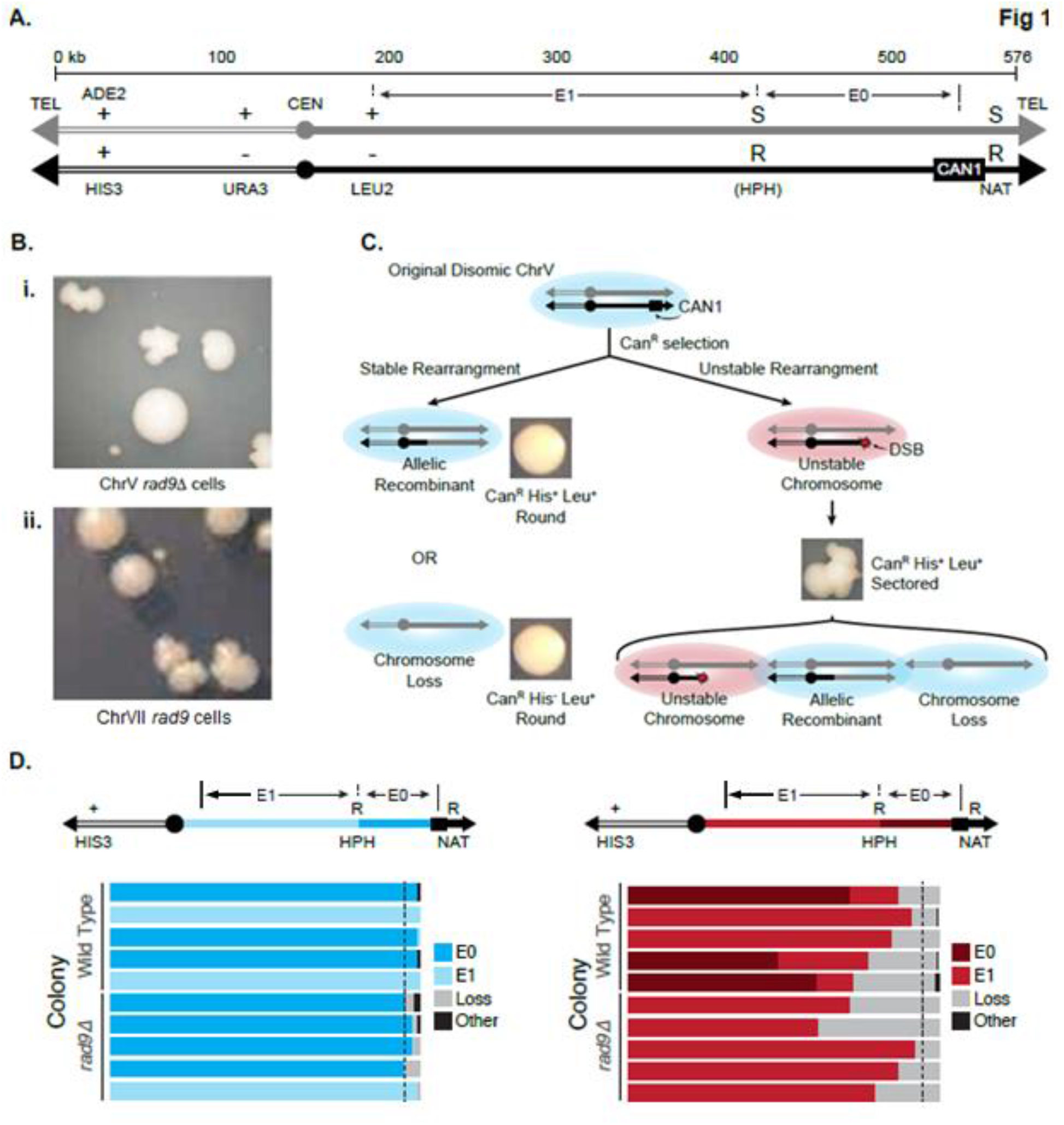
Characterization of the ChrV disome instability system. (A) Schematic of the ChrV disome system. The two homologues of ChrV are shown in black and grey. Rearrangements along the black chromosome are selected by loss of the CAN1 gene. The other heterozygous markers are used to identify the general location of the rearrangement (B) Example of the colony phenotypes in *rad9* mutant strains (Rad9 is a checkpoint protein and a mediator of DNA damage) in the ChrV (i) and ChrVII (ii) disome systems. The colony is either circular, which indicated that the founding cell contained stable chromosomes, or the colony is “sectored,” which indicates that the founding cell contained an unstable chromosome. The colony phenotypes are similar between the ChrV and ChrVII system. (differences in hue of actual colony images are due to different cameras being used, ii was adapted from Beyer and Weinert 2016). (C) Model of the cells fate after Can^R^ selection. Cells can lose CAN1 through a stable rearrangement (allelic recombination or chromosome loss), which results in a round colony. Or cells could undergo an unstable rearrangement that results in a colony with multiple genotypes and a sectored phenotype. (D) Proportion of recombinants from the lineage assay. [Left] Analysis of 10 individual stable, allelic recombinant colonies. Cells were taken from 10 stable allelic recombinant colonies (5 from the wild type strain and 5 from a *rad9Δ* mutant strain) and at least 100 cells were analyzed for the retention of HPH by selection on media plates containing hygromycin. Dashed line indicates 95%. [Right] Analysis of 10 individual unstable sectored colonies. Experiment was conducted as before in Admire et al., 2006, Langston et al., 2020.

Since the number of proteins involved in maintaining the fidelity and replication of chromosomes are vast, we have organized our results by first discussing mutations in genes connected to the biology of three regions of the chromosome; 1) centromere/kinetochore (*mad2Δ* and *pds1Δ*; spindle checkpoint deficient strains), 2) telomere (*cdc13* and *tel1Δ*; telomere maintenance deficient strains), and 3) chromosome-wide (*rad9Δ* and *rad17Δ*; DNA damage checkpoint/repair deficient strains) (Fig. 2). We then investigate mutations in other facets of the DNA repair pathways and their effect on chromosome instability.

**Figure 2.**
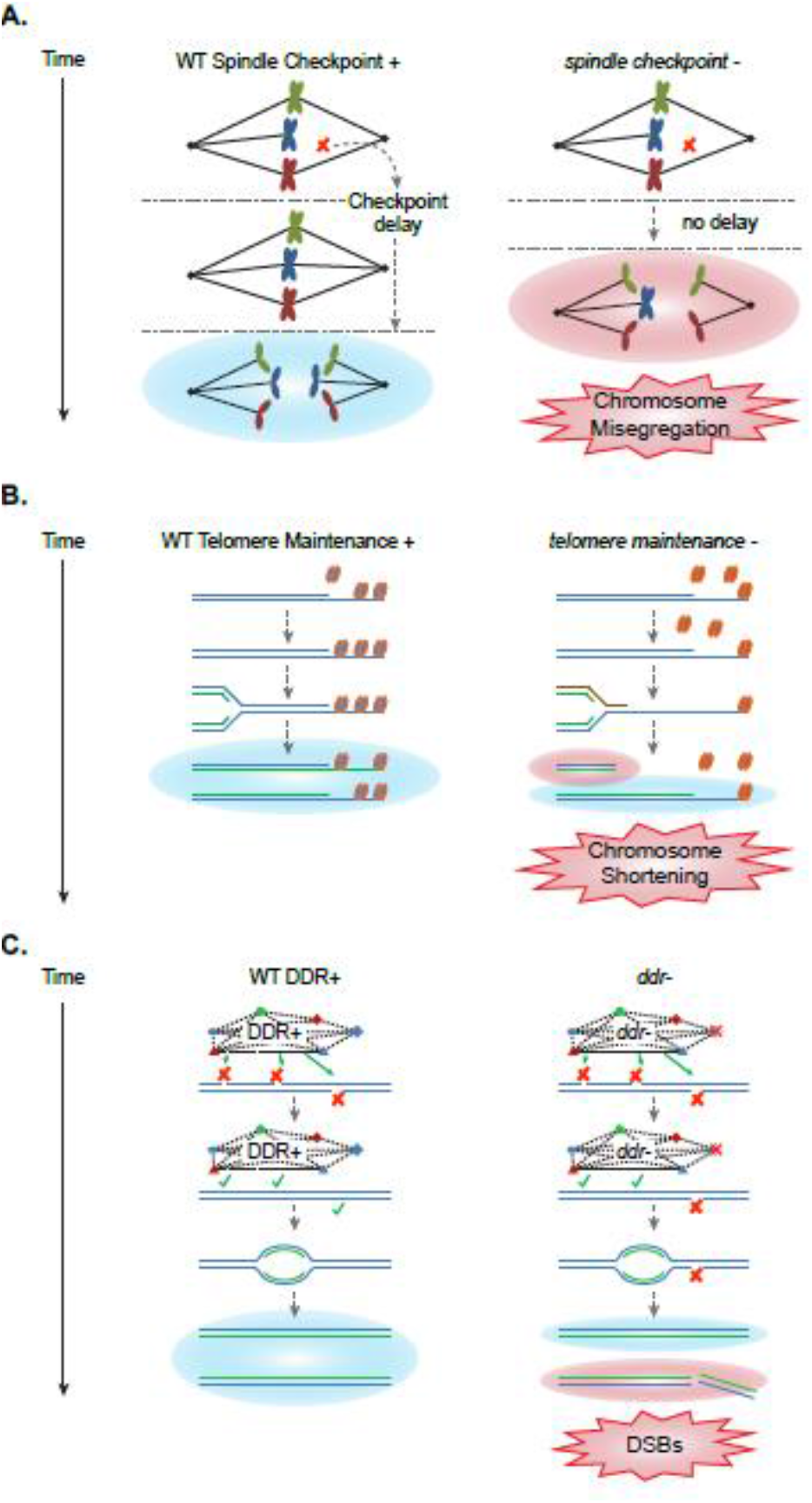
Simplified time lapse (from top to bottom) illustrations of: A) functional Spindle Checkpoint+ (left) ensuring proper chromosome segregation and deficient *spindle checkpoint-*(right) causing chromosome misegregation *(Blue lines represent spindles, green blue and maroon lobes represent replicated chromosomes, red X represents no spindle attachment, green check represent proper spindle attachment)* B) functional Telomere Maintenance+ function (left) ensuring proper telomere protection and deficient *telomere maintenance-*(right) causing chromosome end degradation *(Double blue lines represent double stranded chromosomal DNA, green lines represents newly replicated DNA, blue nodules represent functional telomere proteins, red nodules represent impaired telomere proteins)* C) functional DDR+ (left) ensuring proper chromosome repair and replication and deficient *ddr-*(right) causing chromosome breaks resulting in fragmented chromosome after replication *(Double blue lines represent double stranded chromosomal DNA, green lines represent newly replicated DNA, network represents the many proteins involved in DDR,red Xs on the double blue lines represent DNA breaks, red X on the ddr-network represents a faulty DDR protein)*

Furthermore, to provide a valid comparison of the ChrV disome model system to the ChrVII system we have used the same the laboratory resources and methods as in the previous ChrVII studies (Admire et al., 2006; Paek et al., 2009; Kaochar et al., 2010), i.e., assays, statistical significance calculations, reagents, and growth media.

## MATERIALS AND METHODS

### Yeast strains

The original ChrV disome strain (TY699) is MATa *ade2-1 leu2-3 trp1-1 ura3-52/*URA3, *his3::his3-11,15*/CAN1-NAT GAL psi+ 187520-187620bp::LEU2/+ *can1Δ*::ADE2 *can1Δ*::/HIS3. The ChrV disomic strain was derived from W303a and was originally a gift from Angelica Amon. The strain was modified by Alison Adams and Weinert Labs using the appropriate PCR-amplified markers for transformation followed by, selection and confirmation of the ChrV disome mutants. First, the Adams Lab placed the ADE2 and HIS3 genes to replace the CAN1 loci on the two ChrV homologs. and generated a CAN1:NAT gene pair, and inserted this into site 541300, about 40kb from the right telomere (forming TY800) (A. P. Keller Master Thesis, Northern Arizona University, 2013). Second, the Weinert Lab inserted the wildtype URA3 gene on the “non-CAN1” homolog and the HPH gene into the COM2 ORF, generating TY813. The ChrVII disome used in this study was as reported previously Admire et al., 2006 where mutant strains were created by deleting specific genes with the KanMX allele derived from the SGD strain library.

All ChrV disome mutant strains were generated from TY800 or TY813 by LiAC/ssDNA/PEG transformation using DNA cassettes amplified by polymerase chain reaction (PCR) (Table S1). Unless otherwise stated two isolates for each mutation in ChrV were analyzed for instability.

### Chromosome instability assays

Analyses of instability of the ChrV disome was carried out essentially as described previously for the ChrVII disome (Admire et al. 2006). Briefly, single cells containing both intact ChrV homologs were plated on rich media plates (YEPD, 2% dextrose) and grown for 2-3 days at 30°C to form colonies of between 1 x 10^6^ and 2 x 10^6^ cells per colony; spontaneous chromosome changes occur as the initial cell divides on rich media plates. To then identify cells with chromosome changes, the cells from individual colonies were suspended in double distilled water “ddH20”, counted with a hemocytometer, and plated on selective media containing canavanine (60μg/mL), or canavanine histidine-minus leucine-minus, or complete (SC) media. All media with canavanine lacked arginine and serine to enhance sensitivity to canavanine, a toxic arginine analog. CanR colonies form in 2 to 5 days; round colonies form in 2 to 3 days, while sectored colonies grow slowly and require up to 5 days. We then analyzed the CanR colonies for their colony morphology (round or sectored) and their genetic content as described in Results, identifying chromosome loss, allelic recombinants, and unstable chromosomes, using a “lineage analysis” to confirm that sectored colonies arose from an unstable cell, as described below. We report the median/average instability per viable cell plated on selective media (i.e. ‘10 x 10-5’ = 10 cells out of 100,000 had an unstable event where there was loss of CAN1 gene function). Cell viability was calculated by counting the number of cells that had formed a micro-colony on rich media vs. the number of cells which had not, after 1 day of growth (micro-colony with ≤6 cells indicates an inviable cell).

For each mutant strain tested, we determined the frequencies of events from at least six colonies, representing six biological replicas, from each of two independent genetic isolates, unless otherwise noted. We report the median frequency and standard deviations; statistical significance was calculated using the Kruskal-Wallis method (Kruskal & Wallis, 1952).

### Lineage Assay

We employed a lineage assay to determine if a CanR colony arose from an originating cell that had a stable or an unstable chromosome instability event. Generally, a CanR cell that is stable forms a round colony on selective media in which most cells (>95%) in that colony have the same phenotype. In contrast, a CanR cell that has an unstable chromosome forms a sectored colony on selective media in which cells have different phenotypes (>5% heterogeneity). We typically analyze about 100 cells from each CanR Leu+ His+ colony: about 100 cells from each round or sectored colony are plated onto rich media plates, allowed to divide and form about 100 “secondary colonies” whose phenotypes were determined by replica plating to synthetic media plates, each lacking one genetic marker essential amino acid, or media containing a drug (hygromycin). Cells were grown for 2–3 days at 30°C and then assessed for colony growth for each selective plate. From one CanR Leu+ His+ colony, if greater than 95% of the secondary colonies had the same phenotype, we infer the initial CanR cell from that CanR colony had a stable phenotype. In contrast if less than 95% of the secondary colonies had the same phenotype (i.e. more than 5% of the secondary colonies had different phenotypes), we inferred the originating CanR cell for that CanR colony was unstable.

### Pulse Field Gel Electrophoresis (PFGE) and Southern blots

We sought to determine if any of the unstable chromosomes formed a detectably size altered chromosome using pulse field gel electrophoresis. We identified sectored CanR His+ Leu+ colonies, grew cells in 15 ml of complete selective (his-leu-) liquid media to generate a culture that might retain an unstable chromosome, isolated DNA and performed pulsefield gel analyses using conditions that optimize for separation of 300- and 700-kb chromosomes (Iadonato and Gnirke 1996). In a previous ChrVII study, this strategy allowed identification of translocation and subsequent hotspot of instability in the ChrVII disome (Admire et al. 2006). To detect changes to ChrV, we used probes for the ChrV sequence (115867bp – 117277bp) to the URA3 locus that will identify DNA of both alleles on the two homologs (the ura3-52 and URA3+ allele). We used Roche DIG High Prime DNA Labeling and Detection Starter Kit II, Cat. No. 11 585 614 910.

### Data format and interpretation; Table 1

Instability data in Table 1 were calculated and presented the same way as in previous ChrVII disome studies to allow comparison between the ChrV and ChrVII disome systems (Table S2A, B, C, D). WT-Unstable Chromosomes 10 ± 5.4 x 10^−5^ signifies that on average 10 cells out of 10^5^ cells have a unstable ChrV chromosomes and will grow into sectored colonies (on ‘canavanine histidine-minus leucine-minus’ selective media plates) with a standard deviation of 5.4 (i.e., 10 of the cells have undergone GCR where there is a loss of CAN1 gene function). ‘Fold Instability’ of all mutant strains are relative to WT, e.g., *rad9* Unstable Chromosomes is 300 ± 73 x 10^−5^ therefore fold instability relative to WT is 30 (300/10).

**TABLE 1.** ChrV DISOME MUTANT STRAINS

| TABLE 1 |  |  |  |  |  |  |  |
| --- | --- | --- | --- | --- | --- | --- | --- |
| ChrV DISOME MUTANT STRAINS |  |  |  |  |  |  |  |
| | Genotype | Unstable Chr. ($\times 10^{-5}$) | | Allelic Rec. ($\times 10^{-5}$) | | Chr. Loss ($\times 10^{-5}$) | |
|  |  | mean & std. dev. | fold instab. | mean & std. dev. | fold instab. | mean & std. dev. | fold instab. |
| Wildtype | *ChrV RAD+* | 10 ± 5.4 | 1 | 30 ± 26 | 1 | 5.4 ± 4.5 | 1 |
| Telomere | *V cdc13<sup>b</sup> 30°* | 280 ± 56 | **28\*** | 660 ± 320 | **22\*** | 1500 ± 980 | **280\*** |
| NHEJ | *V lig4Δ* | 3.5 ± 1.5 | **0.35\*** | 38 ± 26 | 1.3 | 1.7 ± 1.8 | 0.31 |
| Spindle check pt. | *V mad2Δ* | 24 ± 6.8 | **2.4\*** | 20 ± 11 | 0.67 | 38 ± 11 | **7.0\*** |
| Securin | *V pds1Δ* | 74 ± 64 | **7.4\*** | 22 ± 15 | 0.73 | 140 ± 120 | **26\*** |
| Mediator | *V rad9Δ* | 300 ± 73 | **30\*** | 24 ± 15 | 0.8 | 180 ± 63 | **33\*** |
| Sensors | *V rad17$\Delta^b$* | 1400 ± 330 | **140\*** | 160 ± 97 | **5.3\*** | 600 ± 1300 | **110\*** |
| PRR | *V rad18Δ* | 400 ± 150 | **74\*** | 110 ± 130 | **3.7\*** | 250 ± 440 | **46\*** |
| HR | *V rad52Δ* | 210 ± 52 | **21\*** | 1.5 ± 2.3 | **0.05\*** | 510 ± 540 | **94\*** |
| Helicase | *V rrm3$\Delta^b$* | 7.9 ± 4.8 | **0.79\*** | 49 ± 24 | **1.6\*** | 41 ± 21 | **7.6\*** |
| PIKK | *V tel1Δ* | 32 ± 24 | **3.2\*\*** | 38 ± 26 | 1.3 | 1.7 ± 1.3 | 0.31 |
| MRX | *V xrs2Δ* | 570 ± 190 | **5.7\*** | 33 ± 70 | 1.1 | 220 ± 68 | **41\*** |
| Double mutations | *V rad9Δ lig4Δ* | 450 ± 100 | **45\*** | 43 ± 26 | 1.4 | 450 ± 420 | **83\*** |
| | *V tel1Δ rad17Δ* | 6400 ± 5000 | **640\*** | 4500 ± 4700 | **150\*** | 1300 ± 1300 | **240\*** |
| Statistically significant in boldface type. Kruskal–Wallis test, *P < 0.01, **P < 0.05. |  |  |  |  |  |  |  |
| <sup>b</sup>Sample sizes: *ChrV disome strains*; *mad2Δ* N=3 for two isolates; *rad17Δ* N=3 for three isolates; *rrm3Δ* N=4 for three isolates; *cdc13* N=4 for one isolate |  |  |  |  |  |  |  |

## RESULTS

### Characteristics and details of the Chromosome V system

The ChrVII and ChrV disome systems are conceptually similar (Fig. 1, Fig. S1). Specifically, both disome systems contain a second copy of the indicated chromosome. This allows for GCRs to stably form in the extra chromosome without affecting the viability of the cell.

The extra chromosome (black) contains the CAN1 gene that encodes arginine permease; when intact, confers sensitivity to the drug canavanine (Fig. 1A). CAN1 allows for negative selection of cells in which the ChrV CAN1 gene has been lost, leading to canavanine resistance (CanR) (Admire et al., 2006; Paek et al., 2009). The ChrV disome contains additional genetic markers (ADE2, HIS3, and LEU2) that allow for selection and analyses of GCRs in which these markers have been disrupted or lost. For strain TY813 the HPH gene (which confers resistance to hygromycin) was inserted into the right arm of ChrV to allow detection of recombination within the EO or E1 intervals (Fig. 1A) for the TY814-816, 818 mutant strains. There are three trackable chromosome arrangements in this system: chromosome loss, allelic recombination in either region E0 or E1 (distinguishable in strains containing HPH; Fig. 1A), and the formation of unstable chromosomes (Fig. 1C).

As observed in the ChrVII system, sectored colonies formed in the ChrV system, in which the originating cell has generated progeny of many different genotypes indicating an unstable ChrV within cells (Fig. 1C). This non-uniform growth is most likely the result of cell death in initially unstable CanR cells, as well as the multiplicity of genotypes and associated growth rate differences within the colony. Furthermore, cells that have lost the CAN1 homologue generated round colonies, and cells where allelic recombination resulted in the loss of CAN1 also generated round colonies (Fig. 1C). We distinguish between chromosome loss and allelic recombination by the presence or loss of the HIS3 gene on the left arm of the CAN1 homologue; loss of HIS3 and CAN1 indicate chromosome loss, while the retention of HIS3 and loss of CAN1 indicate an allelic recombination. Though chromosome truncations may also be occurring to account for the CAN1^-^ and HIS3^+^ phenotype, since there were no previous ChrVII studies that make the distinction of CAN1 loss due to truncation vs. CAN1 loss due to other GCRs, a comparative study of this type is not possible.

To confirm that sectored colonies contain multiple genotypes in the ChrV disome, we used a lineage assay as in the previous ChrVII disome studies (Fig. 1D) (see Materials and Methods for lineage assay details). We found that the cells in each round allelic recombinant colony was mostly of one genotype, with at least 95% identity (Fig. 1D left). In contrast, all of the sectored colonies contained at least 2 of the 3 detectable genotypes in this system (Fig. 1D).

### Proteins in spindle assembly checkpoint stabilize the genome

Mad2 and Pds1 are canonically known for their role in the spindle assembly checkpoint, though both may be involved in other aspects of DNA metabolism (Cohen-Fix et al., 1996; Li & Murray, 1991; Yamamoto et al., 1996). Since the *mad2Δ* mutant strain was previously shown to have increased chromosome loss and unstable chromosomes in the ChrVII system we created and tested a similar mutant strain in the ChrV system.

Mad2 ensures that proper spindle assembly occurs prior to chromosome segregation during mitosis (Fig 2A). Mad2 also appears to be involved in DDR and when deleted has been shown to shorten the time cells are arrested when DSBs occur (Dotiwala et al., 2010) We find that *mad2Δ* cells in both disome systems have a moderate level of unstable chromosome formation, and a high frequency of loss, as one would expect from a spindle defect (Table 1, Table S2D).

Pds1 is also active in both the spindle checkpoint and the DNA damage checkpoint downstream of Chk1 (Sanchez et al., 1999). Pds1/securin prevents Esp1-separase from cleaving cohesions until the metaphase to anaphase transition commences (Ciosk et al., 1998; Cohen-Fix, 2000). Pds1 may have other functions during DNA replication (Clarke et al., 1999). We find that *pds1Δ* mutant cells are extremely unstable in both the ChrV and ChrVII systems, 7.4x and 31x increase in unstable chromosomes, respectively (Table 1 and Table S2D).

There also may be another connection between the spindle checkpoint and DDR as it has recently been reported that DNA replication, a potential source of DNA errors, can happen during mitosis even in budding yeast. Perhaps a failure of checkpoint controls for this type of DNA replication results in instability. (Bhowmick et al., 2016; Ivanova et al., 2018; Min et al., 2017);

### Proteins involved in telomere maintenance are important for genome stability

It has recently been found that telomere maintenance is crucial for maintaining genome stability in the ChrVII disome system (Langston et al., 2020). We sought to test whether telomere health was equally crucial in the ChrV system.

We first tested the single-stranded DNA (ssDNA) binding protein Cdc13 (Fig. 2B). Cdc13 is a part of the heterotrimeric complex CST along with Stn1 and Ten1 and is specific to telomeric ssDNA. It has also been termed “telomeric-RPA” because of its structural similarity with RPA (Gao et al., 2007; Garvik et al., 1995). It was recently demonstrated that Cdc13 prevented chromosome instability originating from a replication-based defect at the telomere (Langston et al., 2020). We therefore tested the same *cdc13* mutant, *cdc13*^*F684S*^ in the Chr V disome system. We found that, like in the ChrVII disome, *cdc13*^*F684S*^ has an increased level of all three types of instability (chromosome loss, allelic recombination, and unstable chromosomes; Table 1, Table S2D). We also note that *cdc13*^*F684S*^ is more unstable in the ChrV disome system compared to the ChrVII disome system.

We also investigated the role of Tel1, the mammalian ATM ortholog and a PIKK protein kinase that responds to double-strand breaks (DSBs) and regulates telomere length (Harrison & Haber, 2006; Morrow et al., 1995). It was previously published that Tel1 plays a role in maintaining chromosome stability in the ChrVII disome system (Kaochar et al., 2010). As expected, *tel1Δ* cells in the ChrV system also exhibit increased levels of all three types of instability (chromosome loss, allelic recombination, and unstable chromosomes; Table 1, Table S2B). We note that, unlike *cdc13*^*F684S*^, instability of *tel1Δ* cells in the ChrV system tended to have less overall chromosome instability than what was found with the ChrVII system.

### The DDR pathway is also crucial for maintaining chromosome stability

Rad9 is a checkpoint protein and a mediator of DNA damage signaling whose functional ortholog is the mammalian 53BP protein (Harrison & Haber, 2006; Weinert & Hartwell, 1993). It was one of the first proteins tested with the ChrVII disome system, therefore we tested *rad9Δ* in the ChrV system and confirmed that *rad9Δ* resulted in an increase in all three types of chromosome instability (chromosome loss, allelic recombination, and unstable chromosomes) (Fig. 2C). This suggests that Rad9 also plays a role in maintaining chromosome stability of ChrV as well as ChrVII (Table 1, Table S2A).

Additionally, we tested the role of Rad17 in maintaining ChrV chromosome stability. Rad17 is a protein of the heterotrimeric checkpoint sliding clamp, a structural orthologue to PCNA that may bind the 5’ recessed end in the lagging strand (Paulovich et al., 1997; Weinert & Hartwell, 1993). In the ChrVII system, *rad17Δ* had an exceptionally high frequency of instability, particularly in unstable chromosomes (Table S2B). In ChrV the trend of high instability continued (Table 1). We note that frequency of instability tended to be higher in the Chr V system than the ChrVII system for both *rad9Δ* and *rad17Δ*. We conclude that defects in Rad9 and Rad17 also destabilized ChrV, probably through the same mechanism as in ChrVII.

We also analyzed a *rad17Δ tel1Δ* double mutation, as it was found that this double mutation is extremely unstable in the ChrVII disome (T. Beyer and T. Weinert, unpublished). Indeed, we find that *rad17Δ tel1Δ* double mutant cells in the ChrV disome system are also extremely unstable, with 1 in 6 cells having chromosomal rearrangements that lose CAN1 (Table 1).

### Unstable chromosomes are not formed through either homologous recombination or non-homologous end-joining

One surprising finding in the ChrVII system was that unstable chromosomes formed without HR or NHEJ (Table S2A). Specifically, formation of unstable chromosomes is independent of Rad52 (HR) and Lig4 (NHEJ). We tested both these pathways in the ChrV disome to elucidate whether this was also true in ChrV, or if there was a particular structure in ChrVII that favored another mechanism.

When we deleted Rad52 in ChrV, we found that the frequency of unstable chromosomes increased, which complements the ChrVII result (Table 1, Table S2A). Additionally, we found that there were virtually no allelic recombinants in the ChrV *rad52Δ* mutant strain, consistent with the removal of HR. The lack of allelic recombinants in *rad52Δ* was also seen in the ChrVII disome system (Table S2A).

Lig4 acts in NHEJ and is an ortholog of the human Lig4. We found that mutant *lig4Δ* cells change little from the wildtype phenotype, in either the ChrV or ChrVII disome system. Furthermore, we asked if unstable chromosomes (in *rad9Δ*) might require NHEJ, a reasonable hypothesis if dicentric chromosomes were formed by fusion between sisters (Pobiega & Marcand, 2010). We find that *rad9Δ lig4Δ* mutant cells form unstable chromosomes in both systems, indicating the NHEJ is not involved in forming unstable chromosomes (Table 1, Table S2A).

While HR and NHEJ may not play a role in generating unstable chromosomes, DSBs may have a role in unstable chromosome generation. It was found that *xrs2Δ* mutant cells are extremely unstable in the ChrVII system (Table S2B). The ChrV *xrs2Δ* strain also exhibits extremely high levels of unstable chromosomes (Table 1). Xrs2 is part of a heterotrimeric MRX protein complex that contains Mre11 and Rad50 (Ivanov et al., 1992; Ritchie & Petes, 2000). The MRX complex is orthologous to the mammalian MRN complex. It binds to DSBs and helps to initiate both the repair of the break (usually through HR) and the activation of the DNA damage checkpoint. This suggests that DSBs are an intermediate in the formation of unstable chromosomes, even though the HR and NHEJ pathways are not involved in creating unstable chromosomes.

### Errors in DNA replication form unstable chromosomes

To test the role of DNA replication induced instability in the ChrV system, we focused on the Rrm3 helicase. Rrm3 removes proteins from chromosomes so as to allow movement of the replication fork (Boulé & Zakian, 2006; Schmidt et al., 2002). We find that *rrm3Δ* mutant cells have a slight instability phenotype in both ChrV and ChrVII systems, resulting in chromosome loss (Table 1, Table S2C) and sectored colonies.

### Post-replication repair is also involved in stabilizing the genome

The Rad18 protein is a member of the post replication repair (PRR) pathway that controls both translesion bypass of DNA polymerases as well as a template-switching error-prone pathway (Masuyama et al., 2005). Rad18 is the orthologue of the mammalian protein of the same name and function. The ChrV disome *rad18Δ* mutant cells are extremely unstable, more so than in ChrVII disome system (Table 1, Table S2A). Rad18, and by implication PRR, is required to keep chromosomes stable but is not involved in forming unstable chromosomes.

### Difference in instability frequency

In general, the frequency of instability in the ChrV disome system was higher than in the ChrVII disome system. We therefore thought it would be worthwhile to determine if the frequency difference was a statistically significant between the ChrV and ChrVII disome systems (i.e. ChrV *rad9Δ* vs. ChrVII *rad9*). Here again we focus on the six mutant strains connected to the biology of three regions of the chromosome (Fig. 2). The comparison shows statistically significant differences between mutant telomere maintenance strains and mutant DNA damage checkpoint/repair strains, but curiously both mutant spindle checkpoint strains were statistically similar.

### Conflicting altered chromosome size data

In the ChrVII disome system, the occurrence of GCRs was visually supported by the use of PFGE and Southern Blot (Admire et al., 2006). Approximately 30% of *rad9* unstable colonies contained altered ChrVII sizes, which supported a biological model of an unstable chromosome structure that produced GCRs which generated altered chromosome sizes. We therefore analyzed chromosome sizes in the ChrV disome system to determine if altered ChrV sizes were also generated. As in the ChrVII system, we obtained sectored colonies from the *rad9Δ* strain and the very unstable *rad17Δ* strain (Table 1), grew cells from each sectored colony (CanR His+ Leu+ cells) in liquid media lacking histidine and leucine to select for the retention of any unstable chromosomes that may have resulted in an altered ChrV size. We then purified intact chromosomes from each culture and separated the chromosomes using PFGE (Fig S2), probing for the URA3 sequence present on both ChrV homologs (see Materials and Methods; PFGE and Southern blots). But in stark contrast to ChrVII results, we failed to detect any altered ChrV sizes in any of the 50+ sectored colonies tested (only a subset is shown in Fig S2). But, it is worth noting that since different budding yeast disome systems have exhibited different fitness and growth phenotypes (Dodgson et al., 2016), in a similar manner, it may be possible that different disome systems exhibit different types of GCRs, and therefore altered ChrVII sizes were detected in the ChrVII system and altered ChrV sizes were not detected in the ChrV system.

## DISCUSSION

In this study many of the trends of chromosome instability of the ChrVII disome system were similarly reproduced in the ChrV disome system. Both systems require the spindle checkpoint, telomere maintenance, and DDR to maintain chromosome stability (Fig. 2A, B, C respectively), as well as the PRR pathway. Both systems exhibit an astoundingly high frequency of chromosome rearrangement in the double mutant strain (*tel1Δ rad17Δ*) (1 in 6 cells), and both systems result in three distinct types of chromosome instability: chromosome loss, allelic recombinants, and unstable chromosomes. Furthermore, unstable chromosomes were formed independently of NHEJ and HR; and neither Lig4 (NHEJ) nor Rad52 (HR) were required for the formation of unstable chromosomes in either ChrV or ChrVII (as previously published). And the high variance in instability data is also similar.

But what is novel and puzzling are the differences between the two systems, in particular the absence of detection of altered ChrV sizes in the ChrV disome system, which conflicts with the altered ChrVII sizes detected in the ChrVII disome system. Trivial explanations for this difference are that the altered ChrVII sizes detected is an artifact of the ChrVII system or that the absence of altered ChrV sizes is an artifact of the ChrV system. Another possible explanation is the absence of a preferred ‘recombination site’ on ChrV. On ChrVII, in contrast to ChrV, there is a suspected recombination site where many recombination events occur, in particular, a region labeled as the “403 E2 site” (Admire et al., 2006). A preferred recombination site may be needed in order to generate detectable altered chromosome sizes. One possible experiment to bring insight into this difference would be to insert the “403 E2 site” into both ChrV homologues of the ChrV system and see if altered ChrV sizes are now detectable.

As previously mentioned, the frequency of instability was generally higher in the ChrV system (Table 1, 2, S2A-D). One explanation for this difference could be due to location of genetic markers inserted into the chromosome and the ‘neighboring-gene effect’ (Baryshnikova and Andrews 2012; Ben-Shitrit et al. 2012). Since genes can affect the transcription of other proximal and distal genes, it is possible the inserted genetic markers in ChrV are more adversely affecting genes involved in DNA repair and integrity. An experiment testing genetic marker location vs. instability frequency could provide insight into frequency of instability differences between the two system. In conclusion, the use of two very similar but distinct model systems can be a valuable tool to provide new insights and allow opportunity to develop more encompassing and refined biologically relevant models.

**TABLE 2.**
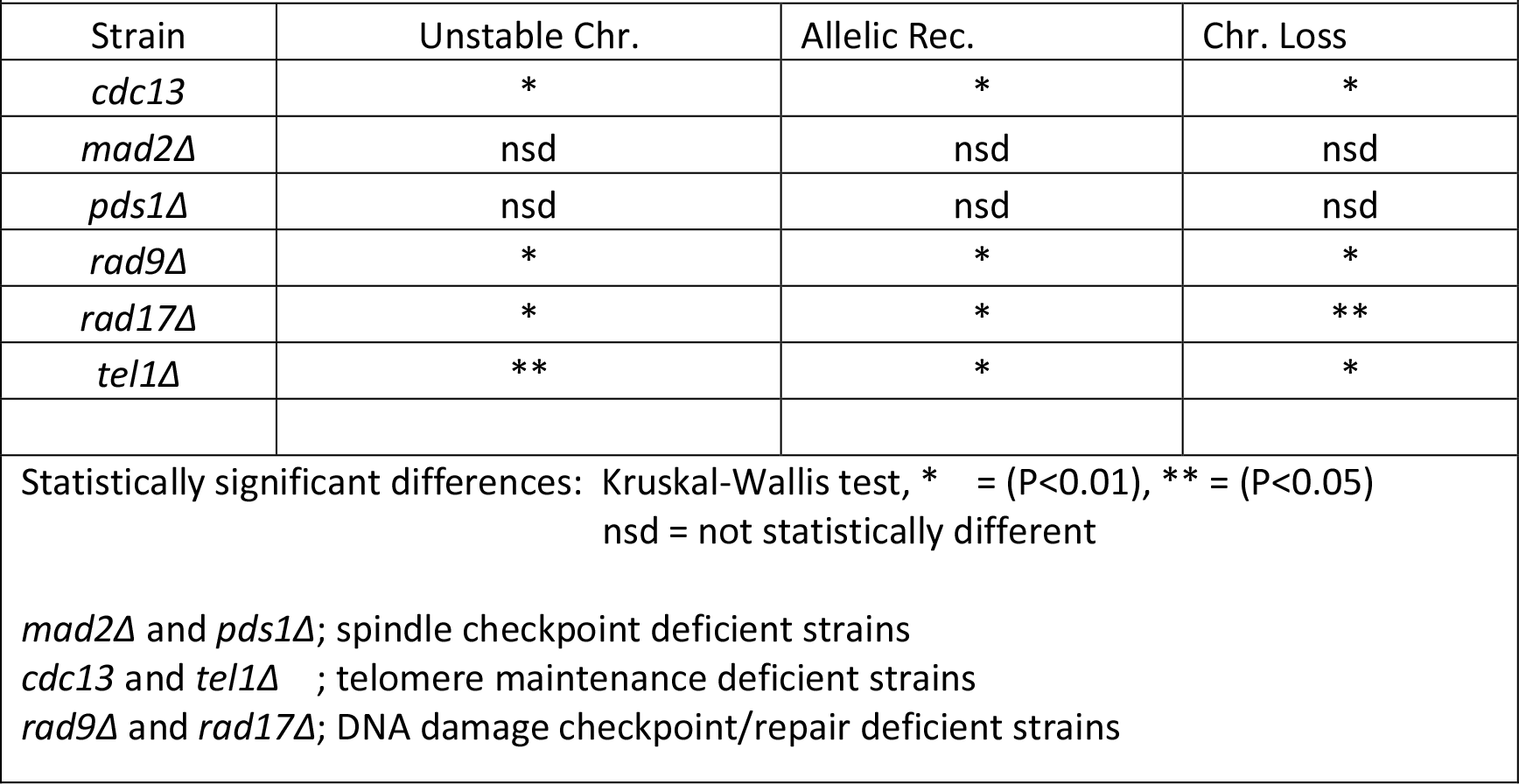
Statistically Significant Differences between ChrV and ChrVII Mutant Strains

## Acknowledgements

*We thank Lisa Shanks and Tracey Beyer for frequent discussions of this work. We also thank Angelika Amon for supplying the 14479 strain used in the development of the ChrV disome system*. *P*.*J*.*V. was funded by T32 GM08659 and self-funded. R*.*E*.*L was funded by T32 GM08659. We declare that no financial or non-financial competing interests exist. Thanks to Chad Park for the use of ChemiDoc for Southern Blot imaging: research reported in this publication was supported by the Office of the Director, National Institutes of Health of the National Institutes of Health under award number S10OD013237*.

## References

Admire, A., Shanks, L., Danzl, N., Wang, M., Weier, U., Stevens, W., Hunt, E., & Weinert, T. (2006). Cycles of chromosome instability are associated with a fragile site and are increased by defects in DNA replication and checkpoint controls in yeast. Genes and Development, 20(2), 159–173. https://doi.org/10.1101/gad.1392506

Baryshnikova, A., and B. Andrews, 2012 Neighboring-gene effect: a genetic uncertainty principle. Nat. Methods 9: 341–343.

Ben-Shitrit, T., N. Yosef, K. Shemesh, R. Sharan, E. Ruppin et al., 2012 Systematic identification of gene annotation errors in the widely used yeast mutation collections. Nat. Methods 9: 373–378.

Beyer, T., & Weinert, T. (2016). Ontogeny of Unstable Chromosomes Generated by Telomere Error in Budding Yeast. PLoS Genetics, 12(10). https://doi.org/10.1371/journal.pgen.1006345

Bhowmick, R., Minocherhomji, S., & Hickson, I. D. (2016). RAD52 Facilitates Mitotic DNA Synthesis Following Replication Stress. Molecular Cell, 64(6), 1117–1126. https://doi.org/10.1016/j.molcel.2016.10.037

Boulé, J. B., & Zakian, V. A. (2006). Roles of Pif1-like helicases in the maintenance of genomic stability. In Nucleic Acids Research (Vol. 34, Issue 15, pp. 4147–4153). https://doi.org/10.1093/nar/gkl561

Breger, K. S., Smith, L., & Thayer, M. J. (2005). Engineering translocations with delayed replication: Evidence for cis control of chromosome replication timing. Human Molecular Genetics, 14(19), 2813–2827. https://doi.org/10.1093/hmg/ddi314

Calabrese, G., Fantasia, D., Franchi, P. G., Morizio, E., Stuppia, L., Gatta, V., Olioso, P., Mingarelli, R., Spadano, A., & Palka, G. (2000). Case report Spectral karyotyping (SKY) refinement of a complex karyotype with t(20;21) in a Ph-positive CML patient submitted to peripheral blood stem cell transplantation. In Bone Marrow Transplantation (Vol. 26). https://www.nature.com/bmt

Chen, C., & Kolodner, R. D. (1999). Gross chromosomal rearrangements in Saccharomyces cerevisiae replication and recombination defective mutants. http://genetics.nature.com

Ciosk, R., Zachariae, W., Michaelis, C., Shevchenko, A., Mann, M., & Nasmyth, K. (1998). An ESP1/PDS1 Complex Regulates Loss of Sister Chromatid Cohesion at the Metaphase to Anaphase Transition in Yeast. In Cell (Vol. 93).

Clarke, D. J., Segal, M., Mondésert, G., & Reed, S. I. (1999). The Pds1 anaphase inhibitor and Mec1 kinase define distinct checkpoints coupling S phase with mitosis in budding yeast A Mec1p-dependent pathway operates early in S phase, but a Pds1p-dependent pathway becomes essential part way through S phase. Results and discussion. http://biomednet.com/elecref/0960982200900365

Cobb, J. A., Schleker, T., Rojas, V., Bjergbaek, L., Tercero, J. A., & Gasser, S. M. (2005). Replisome instability, fork collapse, and gross chromosomal rearrangements arise synergistically from Mec1 kinase and RecQ helicase mutations. Genes and Development, 19(24), 3055–3069. https://doi.org/10.1101/gad.361805

Cohen-Fix, O. (2000). Sister chromatid separation: Falling apart at the seams. Current Biology 2000, Vol 10 No 22, R816–R819

Cohen-Fix, O., Peters, J.-M., Kirschner, M. W., & Koshland, D. (1996). Anaphase initiation in Saccharomyces cerevisiae is controlled by the APC-dependent degradation of the anaphase inhibitor Pdslp. GENES & DEVELOPMENT 10:3081–3093. doi:10.1101/gad.10.24.3081

Dodgson, S. E., Kim, S., Costanzo, M., Baryshnikova, A., Morse, D. L., Kaiser, C. A., Boone, C., & Amon, A. (2016). Chromosome-specific and global effects of aneuploidy in Saccharomyces cerevisiae. Genetics, 202(4), 1395–1409. https://doi.org/10.1534/genetics.115.185660

Dotiwala, F., Harrison, J. C., Jain, S., Sugawara, N., & Haber, J. E. (2010). Mad2 Prolongs DNA Damage Checkpoint Arrest Caused by a Double-Strand Break via a Centromere-Dependent Mechanism. Current Biology, 20(4), 328–332. https://doi.org/10.1016/j.cub.2009.12.033

Gao, H., Cervantes, R. B., Mandell, E. K., Otero, J. H., & Lundblad, V. (2007). RPA-like proteins mediate yeast telomere function. Nature Structural and Molecular Biology, 14(3), 208–214. https://doi.org/10.1038/nsmb1205

Garvik, B., Carson, M., & Hartwell, L. (1995). Single-Stranded DNA Arising at Telomeres in cdc13 Mutants May Constitute a Specific Signal for the RAD9 Checkpoint. In MOLECULAR AND CELLULAR BIOLOGY (Vol. 15, Issue 11). http://mcb.asm.org/

Harrison, J. C., & Haber, J. E. (2006). Surviving the breakup: The DNA damage checkpoint. In Annual Review of Genetics (Vol. 40, pp. 209–235). https://doi.org/10.1146/annurev.genet.40.051206.105231

Hill, H., & Golic, K. G. (2015). Preferential breakpoints in the recovery of broken dicentric chromosomes in drosophila melanogaster. Genetics, 201(2), 563–572. https://doi.org/10.1534/genetics.115.181156

Ivanov, E. L., Korolevt, V. G., & Fabre, F. (1992). Konstantinou Petersburg Nuclear Physics Institute, Academy of Sciences. Genetics 132: 651–664

Ivanova, T., Maier, M., Missarova, A., Ziegler-Birling, C., Carey, L., & Mendoza, M. (2018). Budding yeast complete DNA replication after chromosome segregation begins. BioRxiv, 407957. https://doi.org/10.1101/407957

Iadonato, S.P. and Gnirke, A. 1996. RARE-cleavage analysis of YACs. Methods Mol. Biols. 54: 75–85.

Kaochar, S., Shanks, L., & Weinert, T. (2010). Checkpoint genes and Exo1 regulate nearby inverted repeat fusions that form dicentric chromosomes in Saccharomyces cerevisiae. Proceedings of the National Academy of Sciences of the United States of America, 107(50), 21605–21610. https://doi.org/10.1073/pnas.1001938107

Kruskal, W. H., & Wallis, W. A. (1952). Use of Ranks in One-Criterion Variance Analysis. Journal of the American Statistical Association, 47(260), 583–621. https://doi.org/10.1080/01621459.1952.10483441

Langston, R. E., Palazzola, D., Bonnell, E., Wellinger, R. J., & Weinert, T. (2020). Loss of Cdc13 causes genome instability by a deficiency in replication-dependent telomere capping. PLoS Genetics, 16(4). https://doi.org/10.1371/journal.pgen.1008733

Li, R., & Murray, A. W. (1991). Feedback Control of Mitosis in Budding Yeast. Cell, Vol. 66, 519–531,

Masuyama, S., Tateishi, S., Yomogida, K., Nishimune, Y., Suzuki, K., Sakuraba, Y., Inoue, H., Ogawa, M., & Yamaizumi, M. (2005). Regulated expression and dynamic changes in subnuclear localization of mammalian Rad18 under normal and genotoxic conditions. Genes to Cells, 10(8), 753–762. https://doi.org/10.1111/j.1365-2443.2005.00874.x

Miller, C. R., Stephens, D., Ruppert, A. S., Racke, F., Mcfaddin, A., Breidenbach, H., Lin, H. J., Waller, K., Bannerman, T., Jones, J. A., Woyach, J. A., Andritsos, L. A., Maddocks, K., Zhao, W., Lozanski, G., Flynn, J. M., Grever, M., Byrd, J. C., & Heerema, N. A. (2015). Jumping translocations, a novel finding in chronic lymphocytic leukaemia. British Journal of Haematology, 170(2), 200–207. https://doi.org/10.1111/bjh.13422

Mills, K. D., Ferguson, D. O., & Alt, F. W. (2003). The role of DNA breaks in genomic instability and tumorigenesis. Immunological Reviews Vol. 194: 77–95

Min, J., Wright, W. E., & Shay, J. W. (2017). Alternative Lengthening of Telomeres Mediated by Mitotic DNA Synthesis Engages Break-Induced Replication Processes. https://doi.org/10.1128/MCB

Morrow, D. M., Tagle, D. A., Shiloh, Y., Collins, F. S., & Hieter, P. (1995). Morrow_1995_1-s2.0-0092867495904808-main(2). Cell, 82, 831–840.

Myung, K., & Kolodner, R. D. (2000). Suppression of genome instability by redundant S-phase checkpoint pathways in Saccharomyces cerevisiae. PNAS vol. 99 no. 7, 4500–4507. https://www.pnas.org/cgi/doi10.1073pnas.062702199

Paek, A. L., Kaochar, S., Jones, H., Elezaby, A., Shanks, L., & Weinert, T. (2009). Fusion of nearby inverted repeats by a replication-based mechanism leads to formation of dicentric and acentric chromosomes that cause genome instability in budding yeast. Genes and Development, 23(24), 2861–2875. https://doi.org/10.1101/gad.1862709

Pan, Y., Kytölä, S., Farnebo, F., Wang, N., Lui, W. O., Nupponen, N., Isola, J., Visakorpi, T., Bergerheim, U. S. R., & Larsson, C. (1999). Characterization of chromosomal abnormalities in prostate cancer cell lines by spectral karyotyping. In Cytogenet Cell Genet (Vol. 87). https://www.karger.comwww.karger.com/journals/ccg

Paulovich, A. G., Margulies, R. U., Garvik, B. M., & Hartwell, L. H. (1997). 7, and RAD24 Are Required for S Phase Regulation in Saccharomyces cerevisiae in Response to DNA Damage. Genetics, 145, 45–62

Ritchie, K. B., & Petes, T. D. (2000). The Mre11p/Rad50p/Xrs2p Complex and the Tel1p Function in a Single Pathway for Telomere Maintenance in Yeast. Genetics, 155, 475–479

Rogatto, S. R., Rainho, C. A., Zhang, Z. M., Figueiredo, F., Barbieri-Neto, J., Georgetto, S. M., & Squire, J. A. (1999). Hemangioendothelioma of Bone in a Patient with a Constitutional Supernumerary Marker. https://doi.org/10.1016/S0165-4608(98)00192-7

Sanchez, Y., Bachant, J., Wang, H., Hu, F., Liu, D., Tetzlaff, M., & Elledge, S. J. (1999). Sanchez_et_al_1999_Chk1_Rad53_1166.full(1). Science, 286, 1166–1171.

Schmidt, K. H., Derry, K. L., & Kolodner, R. D. (2002). Saccharomyces cerevisiae RRM3, a 5′ to 3′ DNA helicase, physically interacts with proliferating cell nuclear antigen. Journal of Biological Chemistry, 277(47), 45331–45337. https://doi.org/10.1074/jbc.M207263200

Vasan, S., Deem, A., Ramakrishnan, S., Argueso, J. L., & Malkova, A. (2014). Cascades of Genetic Instability Resulting from Compromised Break-Induced Replication. PLoS Genetics, 10(2). https://doi.org/10.1371/journal.pgen.1004119

Weinert, T. A., & Hartwell, L. H. (1993). Cell Cycle Arrest of cdc Mutants and Specificity of the RAD9 Checkpoint. Genetics, 134, 63–80

Yamamoto, A., Guacci, V., Koshland, D., Guacci, V., Yamamoto, A., Strunnikov, A., Kingsbury, J., Hogan, E., & Meluh, P. (1996). Pdslp Is Required for Faithful Execution of Anaphase in the Yeast, Saccharomyces cerevisiae. The Journal of Cell Biology, Volume 133, Number 1, 85–97

