## Supplemental Tables and Figures for "Comparison of Two Budding Yeast Disome Model Systems: Similarities, Difference, and Conflict"

| **Table S1. *Saccharomyces cerevisiae* strains used in this study** | | |
|---|---|---|
| **ChrV Disome Strains** | | |
| Strain | Genotype[a] | Source |
| TY800 | (Wild Type) | This study |
| TY802 | *rad9Δ::KanMX4* | This study |
| TY804 | *rad17Δ::KanMX4* | This study |
| TY805 | *pds1Δ::KanMX4* | This study |
| TY806 | *mad2Δ::KanMX4* | This study |
| TY807 | *lig4Δ::KanMX4* | This study |
| TY808 | *rad18Δ::KanMX4* | This study |
| TY809 | *rad52Δ::KanMX4* | This study |
| TY810 | *rrm3Δ::KanMX4* | This study |
| TY811 | *tel1Δ::HPH* | This study |
| TY812 | *xrs2Δ::KanMX4* | This study |
| TY813 | *com2::HPH* | This study |
| TY814 | *com2::HPH mad2Δ::KanMX4* | This study |
| TY815 | *com2::HPH rad9Δ::KanMX4* | This study |
| TY816 | *com2::HPH tel1Δ::KanMX4* | This study |
| TY817 | *lig4Δ::KanMX4 rad9Δ::HPH* | This study |
| TY818 | *tel1Δ::HPH rad17Δ::KanMX4* | This study |
| **ChrVII Disome Strains** | | |
| Strain | Genotype[b] | Source |
| TY200 | (Wild Type) | Admire et al 2006 |
| TY206 | *rad9*::ura3 | Admire et al. 2006 |
| TY216 | *rad17::hisGura3* | Admire et al. 2006 |
| TY204 | *rrm3Δ::URA3* | Weinert Lab |
| TY326 | *xrs2Δ::URA3* | Weinert Lab |
| TY362 | *pds1Δ::KanMX4* | Weinert Lab |
| TY440 | *rad18Δ::KanMX4* | Weinert Lab |
| TY629 | *cdc13-F684S*::ura3 (*ts*) | R. Langston and T. Weinert unpublished |
| TY664 | *rad52Δ::KanMX4* | This study |
| TY665 | *mad2Δ::KanMX4* | Vinton and Weinert 2017 |
| TYc27 | *tel1*::*HPH* | This study |
| TYv29 | *lig4Δ::KanMX4* | This study |
| [a]All strains in the ChrV disome section are disomic for ChrV and are derivatives of TY800 MATa *ade2-1 leu2-3 trp1-1 ura3-52/URA3 his3::his3-11,15/CAN1-NAT GAL psi+ can1::ADE2/HIS3, LEU2* replaced 187520-187620bp of the non-*CAN1-NAT* ChrV homolog.<br><br>[b]All strains in the ChrVII disome section are disomic for Chr VII and are derivatives of TY200 MATα *+/hxk2::CAN1 lys5/+ cyh$^r$/CYH$^S$ trp5/+ leu1/+ ade6/+ +/ade3, ura3-2*. | | |

| TABLE S2A | | | | | | | |
|---|---|---|---|---|---|---|---|
| ChrVII DISOME MUTANT STRAINS | | | | | | | |
| | **Genotype** | **Unstable Chr.** (x$10^{-5}$) | | **Allelic Rec.** (x$10^{-5}$) | | **Chr. Loss** (x$10^{-5}$) | |
| | | **mean & std. dev.** | **fold instab.** | **mean & std. dev.** | **fold instab.** | **mean & std. dev.** | **fold instab.** |
| Wildtype | *ChrVII RAD+* | 2.2 ± 1.2 | 1 | 11 ± 12 | 1 | 10 ± 9.2 | 1 |
| NHEJ | *lig4Δ* | 3.2 ± 2.2 | 1.5 | 10 ± 10 | 0.91 | 19 ± 24 | 1.90 |
| Mediator | *rad9Δ* | 50 ± 18 | **23*** | 15 ± 10 | 1.3 | 290 ± 200 | **28*** |
| PRR | *rad18Δ* | 84 ± 33 | **39*** | 7.8 ± 8.9 | **0.7** | 110 ± 150 | **11*** |
| HR | *rad52Δ* | 160 ± 51 | **73*** | < 2.5 | | 23000 ± 10000 | **2200*** |
| Double mutation | *rad9Δ lig4Δ* | 26 ± 6.3 | **12*** | 14 ± 6.8 | 1.2 | 670 ± 780 | 66 |
| Statistically significant in boldface type. Kruskal–Wallis test, *P < 0.05. Data extracted from Paek et al. 2009 | | | | | | | |

| TABLE S2B | | | | | | | |
|---|---|---|---|---|---|---|---|
| **ChrVII DISOME MUTANT STRAINS** | | | | | | | |
| | **Genotype** | **Unstable Chr.** (x$10^{-5}$) | | **Allelic Rec.** (x$10^{-5}$) | | **Chr. Loss** (x$10^{-5}$) | |
| | | **mean & std. dev.** | **fold instab.** | **mean & std. dev.** | **fold instab.** | **mean & std. dev.** | **fold instab.** |
| Wildtype | *ChrVII RAD+* | 3.3 ± 0.8 | 1 | 11 ± 12 | 1 | 13 ± 1.2 | 1 |
| Sensors | *rad17*Δ | 910 ± 290 | **280*** | 160 ± 97 | **5.3*** | 600 ± 1300 | **110*** |
| PIKK | *tel1*Δ | 34 ± 4.2 | **10*** | 6.1 ± 1.3 | 0.6 | 39 ± 18 | 3 |
| MRX | *xrs2*Δ | 340 ± 18 | **100*** | 1.8 ± 0.87 | **0.2*** | 510 ± 88 | **39*** |
| Statistically significant in boldface type. Kruskal–Wallis test, *P < 0.01, data extracted from Kaochar et al. 2010 | | | | | | | |

| TABLE S2C | | | | | | | |
|---|---|---|---|---|---|---|---|
| **ChrVII DISOME MUTANT STRAINS** | | | | | | | |
| | **Genotype** | **Unstable Chr.** (x$10^{-4}$) | | **Allelic Rec.** (x$10^{-4}$) | | **Chr. Loss** (x$10^{-4}$) | |
| | | **mean & std. dev.** | **fold instab.** | **mean & std. dev.** | **fold instab.** | **mean & std. dev.** | **fold instab.** |
| Wildtype | *ChrVII RAD+* | 0.16 ± 0.09 | 1 | 1.3 ± 1.3 | 1 | 5.1 ± 0.6 | 1 |
| Helicase | *rrm3*Δ | 1.3 ± 0.6 | 8.2 | 2.8 ± 2.3 | 2.2 | 8.4 ± 7.8 | 1.6 |
| Data extracted from Admire et al. 2006 | | | | | | | |

| | | | | | | | |
|---|---|---|---|---|---|---|---|
| **TABLE S2D** | | | | | | | |
| **CHRVII DISOME MUTANT STRAINS** | | | | | | | |
| | **Genotype** | **Unstable Chr.** ($x10^{-5}$) | | **Allelic Rec.** ($x10^{-5}$) | | **Chr. Loss** ($x10^{-5}$) | |
| | | **mean & std. dev.** | **fold instab.** | **mean & std. dev.** | **fold instab.** | **mean & std. dev.** | **fold instab.** |
| Wildtype | *ChrVII RAD+* | 5.8 ± 2.8 | 1 | 13 ± 24 | 1 | 7.7 ± 14 | 1 |
| Telomere | *cdc13* 30° | 59 ± 39 | **10*** | 13 ± 13 | 1 | 11 ± 23 | 1.4 |
| Spindle check pt. | *mad2*Δ | 30 ± 24 | **5.2*** | 15 ± 7.9 | 1.2 | 82 ± 87 | **11*** |
| Securin | *pds1*Δ | 180 ± 170 | **31**** | 17 ± 23 | 1.3 | 5700 ± 7500 | **740*** |
| Mediator | *rad9*Δ | 58 ± 34 | **10*** | 6.2 ± 2.9 | 0.5 | 23 ± 13 | **3.0*** |
| Securin | *rad17*Δ | 620 ± 190 | **110*** | 37 ± 37 | 3 | 92 ± 120 | **12**** |
| PIKK | *tel1*Δ | 48 ± 6.1 | **8.3*** | 69 ± 14 | **5.3*** | 38 ± 54 | **4.9*** |
| Statistically significant in boldface type. Kruskal–Wallis test, *P < 0.01, **P < 0.05, data generated in this study. | | | | | | | |

SUPPLEMENTAL FIGURES

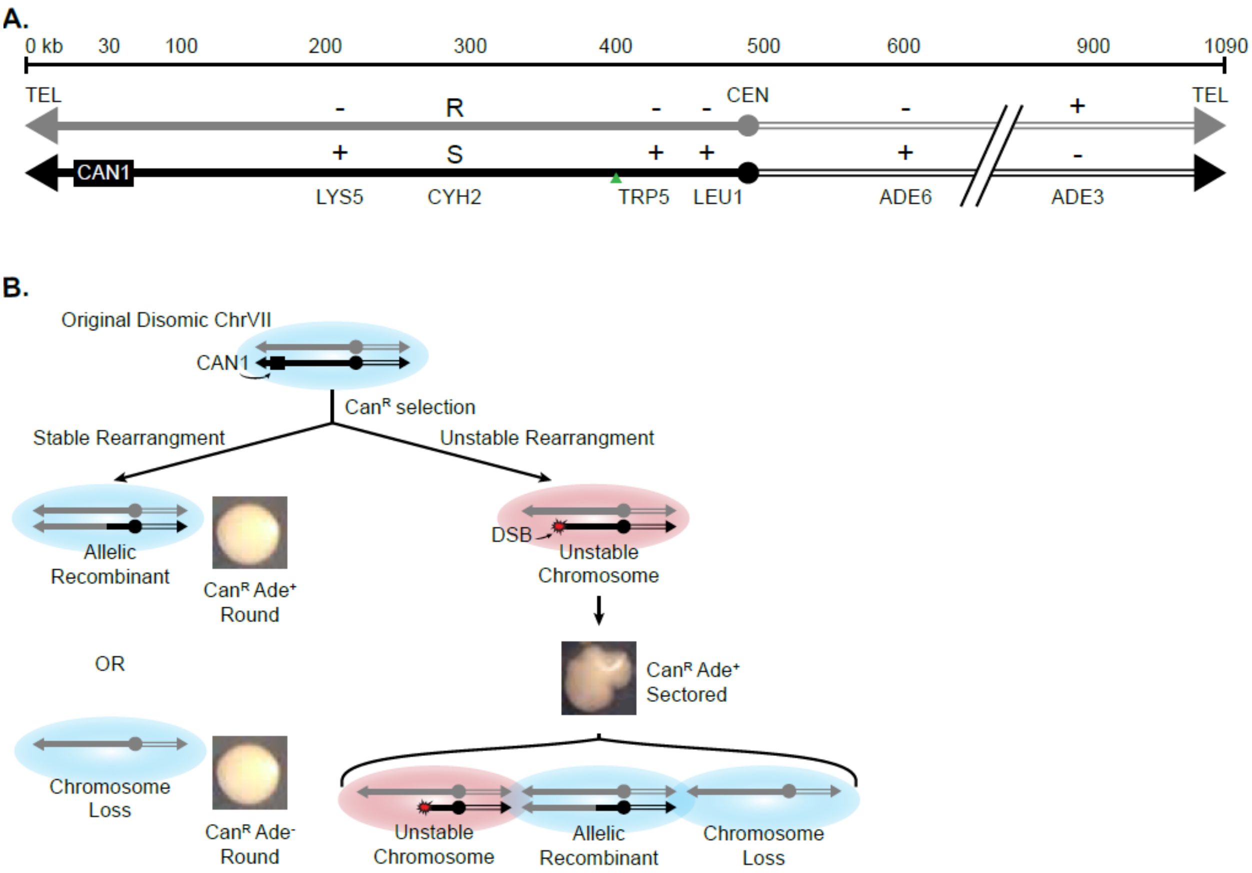

Figure S1. Overview of the ChrVII disome system

(A) Schematic of the ChrVII disome system. The two homologues of ChrVII are shown in black and grey. Rearrangements along the black chromosome are selected for using the CAN1 gene. The other heterozygous markers are used to identify the general location of the rearrangement. The 403 site is marked with the green triangle.

(B) Model of the cells fate after Can$^{R}$ selection. Cells can lose CAN1 through a stable rearrangement (allelic recombination of chromosome loss), which results in a round colony. Or cells could undergo an unstable rearrangement that results in a colony with multiple genotypes and a sectored phenotype.

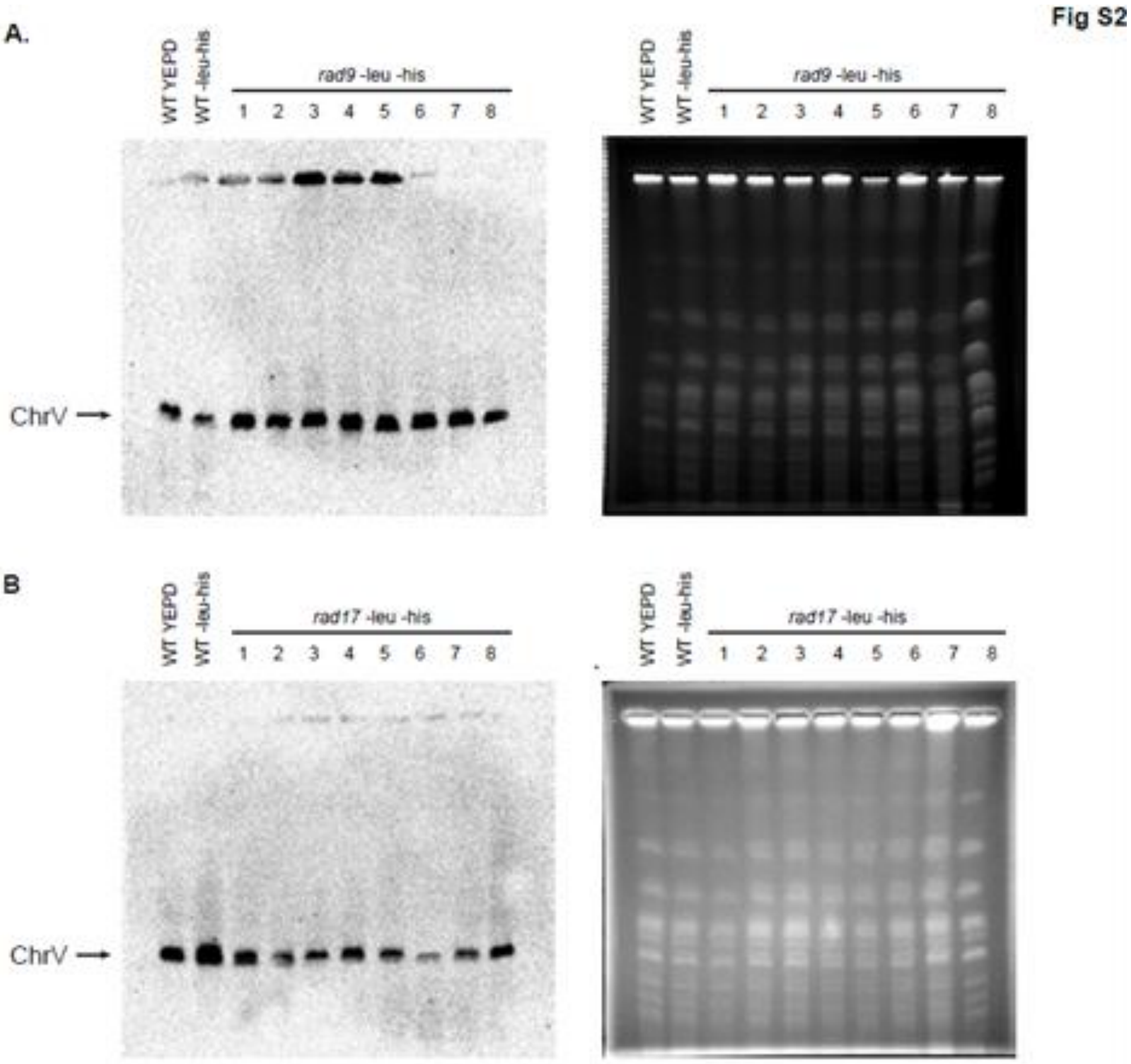

Figure S2. Altered ChrV sizes not detected in *rad9Δ* or *rad17Δ* sectored colonies in the ChrV disome system.

(A) [Left] Southern blot of a PFGE gel of *rad9* sectored colonies using URA3 Dig probes for ChrV. Cells of the indicated genotype were taken from either stock and grown in rich media (without selection) to maintain the original genome or were taken as sectored colonies (with unstable chromosomes) and grown in –leu –his to maintain the rearrangement. 8 individual *rad9* sectored colonies were tested. [Right] PFGE gel used to generate the Southern blot

(B) [Left] Southern blot of a PFGE gel of *rad17* sectored colonies using URA3 Dig probes for ChrV. Cells were prepared as in Fig S1A. [Right] PFGE gel used to generate the Southern blot
